# Evolution along allometric lines of least resistance: Morphological differentiation in *Pristurus* geckos

**DOI:** 10.1101/2022.11.28.518148

**Authors:** Héctor Tejero-Cicuéndez, Iris Menéndez, Adrián Talavera, Gabriel Riaño, Bernat Burriel-Carranza, Marc Simó-Riudalbas, Salvador Carranza, Dean C. Adams

## Abstract

Species living in distinct habitats often experience unique ecological selective pressures, which can drive phenotypic divergence. However, how ecophenotypic patterns are affected by allometric trends and trait integration levels is less well understood. Here we evaluate the role of allometry in shaping body size and shape diversity in *Pristurus* geckos utilizing differing habitats. We found that patterns of body shape allometry and integration were distinct in species with different habitat preferences, with ground-dwelling *Pristurus* displaying the most divergent allometric trend and the strongest integration. There was also strong concordance between static allometry across individuals and evolutionary allometry among species, revealing that body shape differences among individuals were predictive of evolutionary changes across the phylogeny at macroevolutionary scales. This suggested that phenotypic evolution occurred along allometric lines of least resistance, with allometric trajectories imposing a strong influence on the magnitude and direction of size and shape changes across the phylogeny. When viewed in phylomorphospace, the largest rock-dwelling species were most similar in body shape to the smallest ground-dwelling species, and vice versa. Thus, in *Pristurus*, phenotypic evolution along the differing habitat-based allometric trajectories resulted in similar body shapes at differing body sizes in distinct ecological habitats.

## 1. Introduction

Understanding how phenotypic diversity evolves, and elucidating the forces that generate and maintain this diversity, are major goals in evolutionary biology. Because adaptive evolution is the product of natural selection, changes in ecological selection pressures are expected to affect the evolutionary trajectory of phenotypic traits that facilitate an organism’s survival in their habitat. Evolutionary theory predicts that differing habitats will exert unique ecological selection pressures on organisms, resulting in associations between ecological and phenotypic traits. Indeed, species inhabiting differing habitats often display functional, behavioral, or phenotypic differences, that have presumably been the result of adaptive diversification in their respective ecological contexts [1–5].

One possible evolutionary outcome of ecological specialization is that organisms inhabiting similar environments display common phenotypic characteristics. When such patterns occur repeatedly [6,7], this convergent evolution is treated as strong evidence of adaptation. Indeed the ecomorphological paradigm [8] is predicated, in part, on such cases, which emphasize the strong association between the phenotypic traits that organisms display (morphological, behavioral, or physiological) and the ecological characteristics of their habitat that mediate organismal performance. In vertebrates, ecomorphological trends have been well studied in numerous taxonomic groups, and include the emblematic ‘ecomorphs’ of Caribbean *Anolis* lizards that exploit different microhabitats [6,9,10], differential beak morphology in species of Darwin’s finches [11–13], the recurring phenotypes of African lake cichlids across ecological regimes [14,15], and the distinct body forms of freshwater fishes in benthic and limnetic habitats [16–18], among others.

However, while the patterns of morphological differences in distinct ecological contexts have been well documented, less-well understood is how this differentiation has been influenced by trait covariation associated with body size differences (i.e., allometry). Evaluating allometric trends across hierarchical levels (e.g., comparing allometry at the individual level, or static allometry, and among species, or evolutionary allometry) may aid in our understanding of how adaptive morphological change occurs at macroevolutionary scales [19]. It has long been recognized that the interrelationships among traits can exert a strong influence on how phenotypic evolution proceeds, as trait correlations influence the degree to which phenotypic variation is exposed to selection [20]. Thus, the integration among traits can constrain phenotypic change in certain directions, or enhance variation along other phenotypic axes [20–27]. Further, because nearly all linear traits covary strongly with overall body size [28,29], allometric trends could be considered the quintessential expression of phenotypic integration. Thus, identifying whether allometric patterns differ across habitats, and how such patterns of trait covariation affect ecomorphological trends among species utilizing those habitats, remains an important question worthy of investigation.

The Afro-Arabian geckos in the genus *Pristurus* afford the opportunity to elucidate the inter-digitating effects of allometry and habitat specialization on clade-level patterns of phenotypic diversity. Prior work on this system [30] revealed that the colonization of ground habitats has been a trigger of morphological change, specifically reflected in an increase in body size and shape disparity. Interestingly, some ground-dwelling species are among the largest of the genus and also show increased relative head sizes and limb proportions, while some other species with this ecological specialization have evolved to be among the smallest of the group. Additionally, among the species exploiting rocky habitats (the most common ecological feature in *Pristurus),* there are also species with both considerably large and small body sizes [30]. What remains unexplored, however, is how the evolution of body shape is related to differences in body size and whether habitat specialization has an impact in this shape-size relationship.

In this study, we employed a combination of multivariate morphometric and phylogenetic comparative analyses to interrogate macroevolutionary patterns of evolutionary allometry in *Pristurus* geckos of Afro-Arabia. Using phenotypic, phylogenetic, and ecological data, we first characterized allometric trends in body form in the group, to discern the extent to which evolutionary allometric trends across the phylogeny aligned with habitat-based static allometry for species occupying distinct ecological regimes. We then examined changes in allometric trends across the phylogeny, and linked these patterns to overall phenotypic integration, diversification in morphospace, and habitat utilization among taxa. Our analyses reveal that patterns of evolutionary allometry across species align with allometric trends within habitats, demonstrating that the interplay between ecological specialization and allometric trajectories in species with disparate body size may play a determinant role in shaping the phenotypic evolution and hence in adaptive dynamics in this clade.

## 2. Materials and Methods

### (a) Data

We used a combination of phenotypic, phylogenetic, and ecological data to characterize and evaluate intra- and interspecific allometric trends. The data utilized here were obtained from our prior work on this system [30,31], and are briefly described here. First we used a time-dated, molecular phylogeny of squamates that included all members of the genus *Pristurus,* including several currently undescribed taxa. The tree was estimated in a Bayesian framework, using five mitochondrial markers, six nuclear markers, and 21 calibration points [31]. Next we categorized each species as belonging to one of three ecological groups (ground, rock, or tree), based on descriptions of habitat use found in the literature [30]. Finally, we obtained a phenotypic data set containing body size (snout-vent length: SVL) and eight linear measurements (Figure 1) that described overall body form: trunk length (TL), head length (HL), head width (HW), head height (HH), humerus length (Lhu), ulna length (Lun), femur length (Lfe), and tibia length (Ltb) [30]. We restricted our study to those species represented by nine or more individuals; resulting in a dataset of 687 individuals from 25 species (invidivuals per species: *μ* = 27; min = 9, max = 56). Species in the phenotypic dataset were then matched to the phylogeny, which was subsequently pruned to the final topology. All measurements were log-transformed prior to statistical analyses. Additional details regarding data collection and formal descriptions of each linear measurement may be found in the original sources [30,31]. The data are available on DRYAD: https://doi.org/10.5061/dryad.xwdbrv1f6 [32].

**Figure 1.**
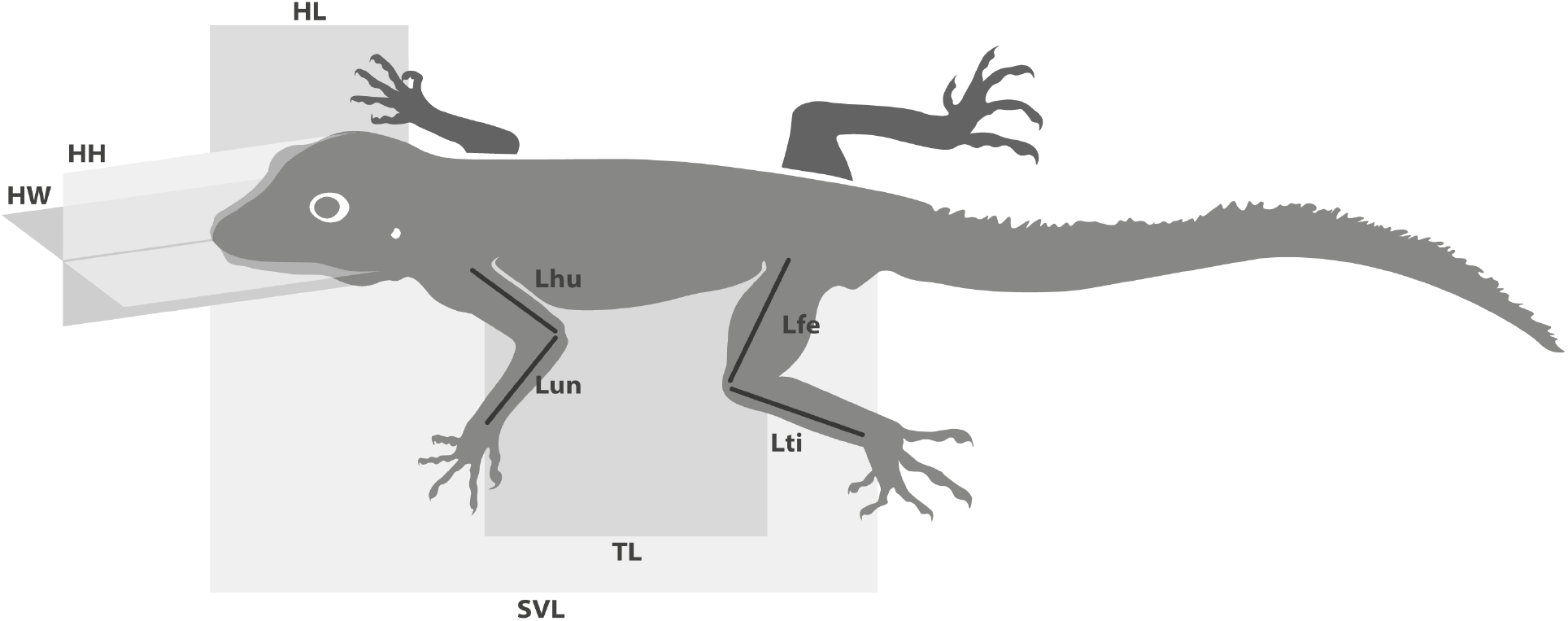
Linear Measurements used in this study. SVL = snout-vent length, TL = trunk length, HL = head length, HW = head width, HH = head height, Lhu = humerus length, Lun = ulna length, Lfe = femur length, Ltb = tibia length [30].

### (b) Statistical and Comparative Analyses

We conducted a series of analyses to interrogate allometric trends, patterns of integration, and macroevolutionary changes in allometry, relative to differentiation in body form. First we characterized evolutionary allometry in the genus by performing a phylogenetic multivariate regression of body form on body size (i.e., SVL), using the species means as data. We then performed an analogous procedure at the individual level, regressing body form on body size using our entire dataset. From both the species-level (phylogenetic) and the individual-level analyses, we obtained the set of regression coefficients, and calculated the difference in their angular direction to describe the extent to which patterns of allometry at the individual level were concordant with evolutionary allometric trends across species.

Next we used the dataset containing all individuals to determine whether trends in static allometry differed across habitat groups. This was accomplished by performing a multivariate analysis of covariance, with body size (*SVL*), *habitat,* and *SVL* × *habitat* as model effects. Significance was evaluated using 999 iterations of a permutation procedure, where residuals from a reduced model were randomly permuted in each permutation (RRPP), model statistics were recalculated, and used to generate empirical null sampling distributions to evaluate the observed test statistics [33–35]. We then compared the multivariate allometric vectors for each habitat group to one another, and to a vector representing multivariate isometry, by calculating pairwise differences in their angular direction in morphospace, and evaluating these relative to empirical sampling distributions obtained through RRPP [34,36,37]. Here, residuals were obtained from a common isometry reduced model, whose common slope component described a pattern of multivariate isometry, and whose intercepts allowed for differences in least-squares means among groups. Patterns of multivariate allometry relative to body size were visualized via regression scores [38] and predicted lines [39], based on the coefficients and fitted values from the linear model described above.

Additionally, because allometry describes the extent to which traits covary with body size and with each other (i.e., integration), we conducted an analysis of integration. Here we characterized the extent of morphological integration in body form for individuals within each habitat group by summarizing the dispersion of eigenvalues of their respective trait covariance matrix [40]. This measure (*V_rel_*) was subsequently converted to an effect size (a *Z*-score), which quantified the strength of morphological integration [41]. We then performed a series of two-sample tests to compare the strength of morphological integration across habitat groups, following the procedures of [41]. Additionally and for comparison, we repeated these analyses on the set of size-standardized trait data, found as a set of shape ratios [42] where each trait was divided by body size (Supplementary Material).

To determine the extent to which static and evolutionary allometry were concordant, we evaluated the directions in morphospace of both the evolutionary (species-level) and static (habitat-based) allometric trends. Specifically, we obtained the set of regression coefficients from both the phylogenetic multivariate regression and the multivariate analysis of covariance analyses above, and calculated the differences in angular direction between the evolutionary trajectory and the static allometry trend for each habitat group. The observed angles were then statistically evaluated relative to empirical sampling distributions obtained through permutation (RRPP), based on the common isometry model described above.

Next, to discern how allometric trends resulted in the evolution of distinct body forms, we examined changes in the body shape proportions across the phylogeny. Here we treated the head dimensions and limb dimensions separately, as allometric trends could potentially differ between these body regions due to differential functional or selective constraints [43]. Because both the head and limb data were multivariate, we first performed a partial least squares (PLS) analysis [44] of the head traits versus SVL, and the limb traits versus SVL, to describe the direction of maximal covariation between each body region and size. We then measured the mean residuals of each species to the inferred allometric trend, which described the extent to which head and limb proportions of species were greater or smaller than expected for their body size. The species residuals were then mapped on the phylogeny of *Pristurus* using a Brownian motion model of evolution, to qualitatively evaluate shifts in head and limbs proportionality across the phylogeny for the group. Similarly, within-species patterns of static allometry were visualized by plotting regressions of PLS scores on SVL for both head and limb traits separately.

Finally, to relate within-species allometric trends with patterns of phenotypic diversification in the group we generated a phylomorphospace, based on a phylogenetic principal component analyses (PCA) on the size-standardized species means obtained from a phylogenetic regression [30]. Here, phenotypic similarities among species, relative to their phylogenetic relationships and habitat affiliations, were observed. Additionally, representative specimens (scaled to unit size) were also visually compared to aid in describing these trends. A similar phylomorphospace was constructed for species means not corrected for body size, and the phenotypic disparity among species means in each habitat was calculated and subsequently compared (Supplementary Material). All analyses were conducted in R 4.2.1 [45], using RRPP version 1.3.1 [46,47] and `geomorph` 4.0.4 [48] for statistical analyses and the `tidyverse` version 1.3.0 [49], `phytools` version 0.7-77 [50], and a modified version of the function `ggphylomorpho` [https://github.com/wabarr/ggphylomorpho] for data manipulation and visualization, as well as scripts written by the authors (Supplementary Material).

## 3. Results

Using phylogenetic regression, we found significant evolutionary allometry in body form across species (*N_sp_* = 25; *F* = 217.9; *Z* = 5.53; *P* < 0.001). Likewise, when allometry in body form was examined across individuals, a similar pattern was observed (*N_ind_* = 687; *F* = 7910.8; *Z* = 9.20; *P* < 0.001). Further, the vectors of regression coefficients between the two analyses were highly correlated (*ρ* = 0.94) and were oriented in nearly parallel directions in morphospace (*θ* = 1.49°). This revealed that the pattern of multivariate allometry across individuals was concordant with macroevolutionary trends of interspecific allometry among species of *Pristurus* across the phylogeny.

Our analyses also exposed significant differences in the allometry of body form among *Pristurus* utilizing distinct habitats (Table 1). Further, pairwise comparisons of multivariate allometric vectors revealed that patterns of static allometry in each habitat differed significantly from isometry, indicating the presence of multivariate allometry in each (Table 2). Additionally, comparisons identified that ground-dwelling *Pristurus* displayed the most distinct allometric trend as compared with *Pristurus* occupying both the rock and tree habitats (Table 2; Figure 2). Here, regression coefficients of each trait versus size (Supplementary Material) revealed that ground-dwelling *Pristurus* exhibited strong positive allometry for all head and limb traits (i.e., *β* > 1.0). By contrast, rock and tree-dwelling *Pristurus* displayed negative allometry (i.e., *β* < 1.0) for head traits, and were more varied for limb traits; with rock-dwelling *Pristurus* displaying positive limb allometry (though less extreme than that of ground-dwelling taxa), whereas most limb traits in tree-dwelling taxa showed negative allometry or near-isometry (Supplementary Material). Thus, these findings implied that larger individuals of ground-dwelling *Pristurus* species displayed disproportionately larger heads and limbs, as compared with large individuals in taxa utilizing other habitat types. Multivariate visualizations of these multivariate allometric trends (Figure 2) confirmed these statistical findings, and indicated that the allometric trajectory in ground-dwelling *Pristurus* was more extreme as compared with either rock- or tree-dwelling *Pristurus*.

**Figure 2.**
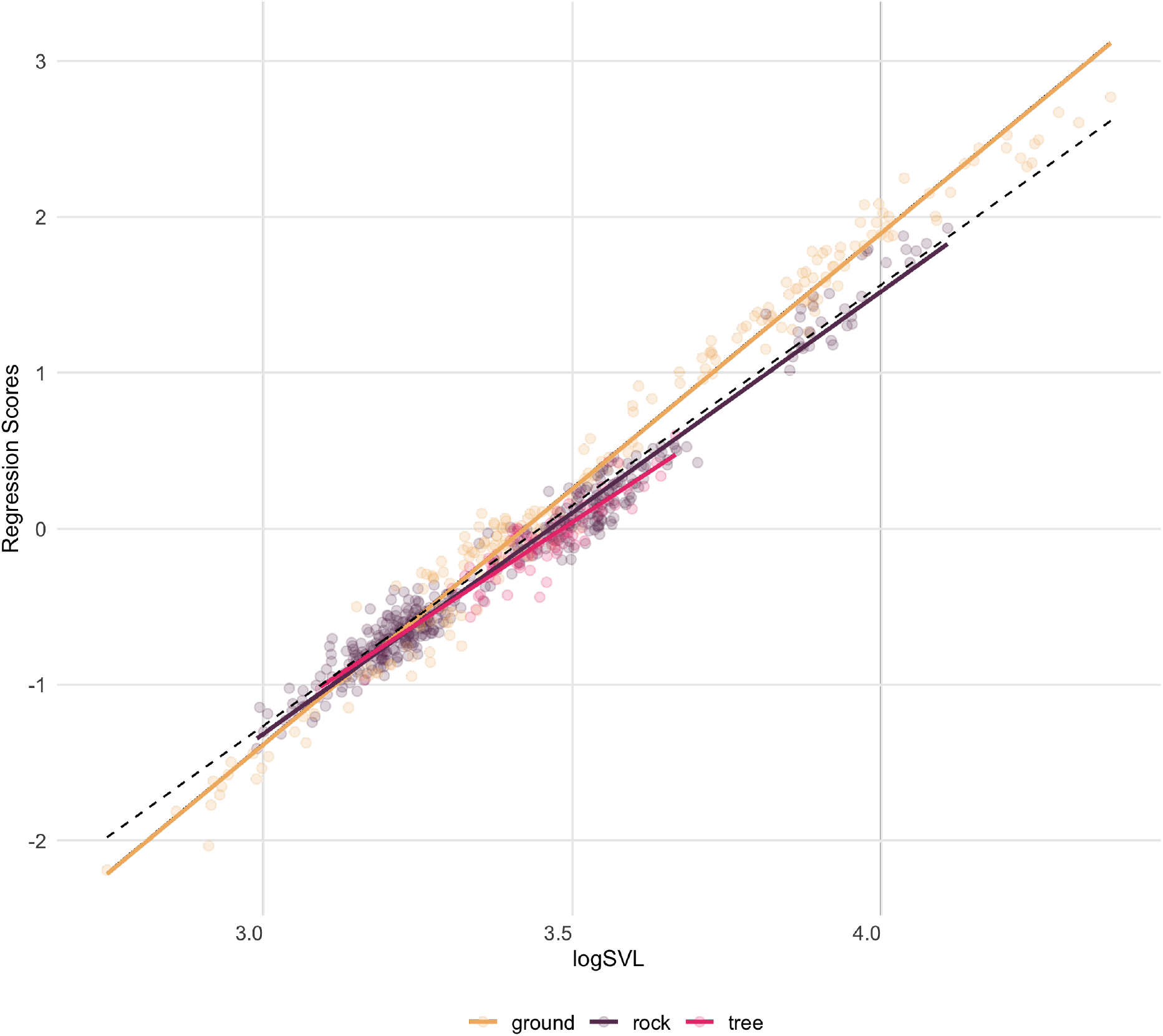
Plot of regression scores and predicted lines representing the relationship between linear body measurements and size (SVL). Individuals are colored by habitat use: ground (beige), rock (dark purple), and tree (magenta). Isometric trend represented by the dashed line.

**Table 1:** Multivariate analysis of covariance describing variation in body form in *Pristurus*.

|  | Df | SS | MS | Rsqr | F | Z | Pr(>F) |
| --- | --- | --- | --- | --- | --- | --- | --- |
| SVL | 1 | 516.04 | 516.04 | 0.92 | 10188.70 | 9.49 | 0.001 |
| habitat | 2 | 6.22 | 3.11 | 0.01 | 61.39 | 9.32 | 0.001 |
| SVL:habitat | 2 | 3.97 | 1.99 | 0.01 | 39.23 | 7.08 | 0.001 |
| Residuals | 681 | 34.49 | 0.05 | 0.06 |  |  |  |
| Total | 686 | 560.72 |  |  |  |  |  |

**Table 2:** Pairwise comparisons of multivariate static allometry for each habitat group. Comparisons with the vector of multivariate isometry are included. Displayed are: pairwise angular differences (*0*_12_), their associated effect sizes (*Z*_*θ*_12__), and significance levels obtained via permutation (RRPP).

|  | Ground | Rock | Tree | Isometry |
| --- | --- | --- | --- | --- |
| **Angle** |  |  |  |  |
| Ground | 0 |  |  |  |
| Rock | 6.629 | 0 |  |  |
| Tree | 8.095 | 3.628 | 0 |  |
| Isometry | 5.034 | 5.901 | 7.189 | 0 |
| **Effect Size** |  |  |  |  |
| Ground | 0 |  |  |  |
| Rock | 7.004 | 0 |  |  |
| Tree | 2.1 | -0.408 | 0 |  |
| Isometry | 7.673 | 7.357 | 1.779 | 0 |
| **P-value** |  |  |  |  |
| Ground | 1 |  |  |  |
| Rock | 0.001 | 1 |  |  |
| Tree | 0.027 | 0.673 | 1 |  |
| Isometry | 0.001 | 0.001 | 0.042 | 1 |

Examination of patterns of trait covariation revealed strong levels of morphological integration within each habitat type (*Z_ground_* = 3.97; *Z_rock_* = 3.72; *Z_tree_* = 2.15). Further, two-sample tests revealed that the strength of morphological integration was significantly greater in ground-dwelling *Pristurus* than either those utilizing rock (*Z_ground-rock_* = 6.59; *P* << 0.001) or tree habitats (*Z_ground–tree_* = 11.17; *P* << 0.001). Arboreal *Pristurus* displayed the lowest levels of integration, which were also significantly lower than in the rock habitat (*Z_rock-tree_* = 7.19; *P* << 0.001). When size was accounted for in the data, levels of integration dropped considerably, though the overall pattern and differences among habitat groups remained the same (Supplementary Material).

Comparisons of evolutionary allometry with static allometry in each habitat revealed substantial concordance between allometric trends at these hierarchical levels. Here, vectors of regression coefficients representing static allometry within habitat groups were oriented in very similar directions with the regression vector representing evolutionary allometry, with small pairwise angles between them (*θ*: 2.3° → 5.9°). Subsequent permutation tests indicated no differences between the static allometry vectors and the regression vector representing evolutionary allometry, indicating strong congruence between them (Table 3). Notably, static allometry in ground-dwelling *Pristurus* was most similar to trends of evolutionary allometry, displaying the smallest angular difference and largest effect size. Thus, static and evolutionary allometry trends were essentially parallel in this group, indicating a direct correspondence between the two. This result implied that phenotypic evolution across species aligned closely with directions of allometric variation within habitat groups at the individual level; namely that larger individuals and larger ground-dwelling species exhibited disproportionately larger heads and limbs, while smaller individuals and smaller ground-dwelling species displayed disproportionately smaller heads and limbs.

**Table 3:** Pairwise comparisons of multivariate evolutionary allometry versus static allometry for each habitat group. Pairwise angular differences between evolutionary and static allometry (*θ_ES_*), their associated effect sizes (*Z_θ_ES__*), and significance levels are displayed.

| | $\theta_{ES}$ | $Z_{\theta_{ES}}$ | P-value |
| --- | --- | --- | --- |
| Evol. vs. Ground | 2.37 | -4.26 | 1.000 |
| Evol. vs. Rock | 4.55 | 0.87 | 0.191 |
| Evol. vs. Tree | 5.96 | 0.21 | 0.405 |

Mapping the residuals of species into the phylogeny showed that large ground-dwelling species displayed greater head proportions than large rock-dwelling species, who exhibited smaller heads relative to body size (Figure 3A). Conversely, the opposite pattern was observed when comparing small species utilizing these habitats: ground-dwelling species showed small relative head proportions while rock-dwelling species displayed generally larger head proportions. In contrast, limb shape showed more variable patterns. Although all large ground-dwelling species consistently displayed large relative limb proportions, large rock-dwelling species were more variable in this trait, with *P. insignis* exhibiting large and *P. insignoides* small limb proportions. For small species, shifts in relative limb proportions seemed more independent of habitat utilization, since there were differences in limb residuals both within rock- and ground-dwelling species (Figure 3B). Visual inspection of static allometry trends within species (Figure 4) largely confirmed these patterns, illustrating that ground-dwelling species generally displayed steeper allometric patterns in head proportions as compared with rock-dwelling species. Overall there was general concordance across taxa in terms of trends of multivariate allometry, affirming that the association between evolutionary allometry and habitat-based static allometry was robust.

**Figure 3.**
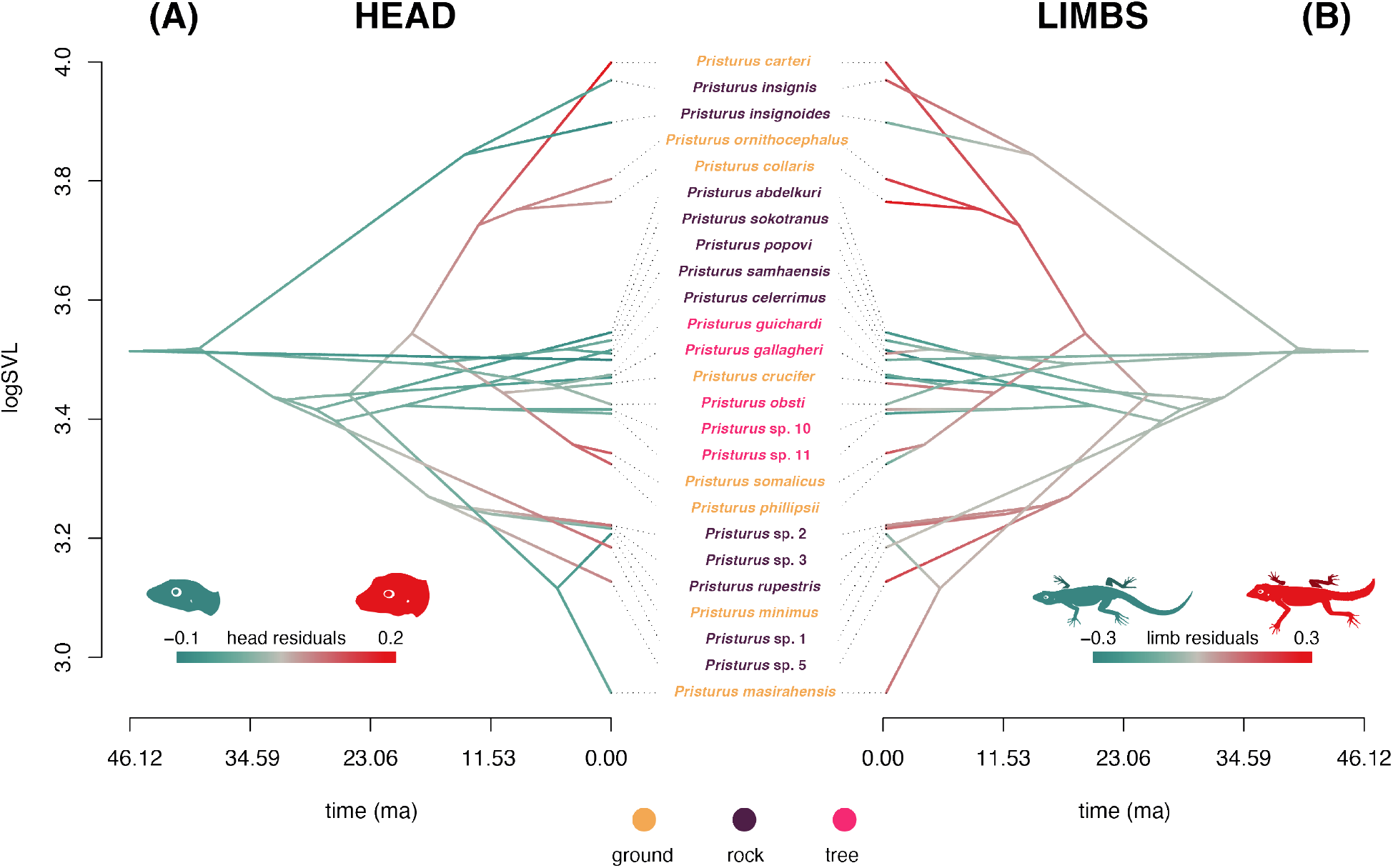
Traitgrams showing the evolution of body size (SVL) through time based on the phylogenetic tree of *Pristurus*. Colors represent an evolutionary mapping of residuals from phylogenetic regressions describing the relationship of (A) head morphology versus body size, and (B) limb proportions versus body size (see text for descriptions). Species names are colored by habitat use: ground (beige), rock (dark purple), and tree (magenta).

**Figure 4.**
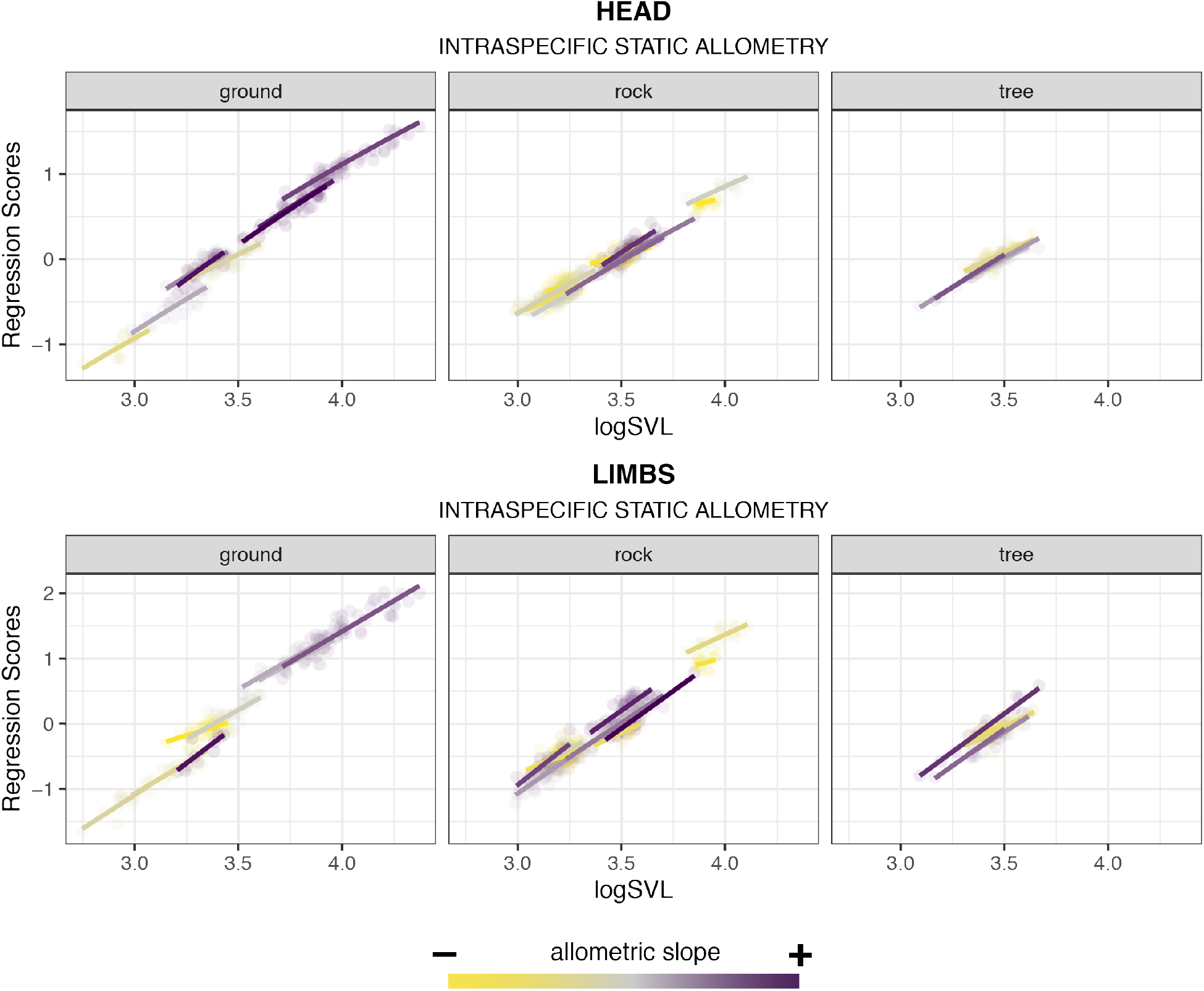
Patterns of static allometry for each species for head traits (upper panel) and limb traits (lower panel). Species are separated by their habitat groups and colored by the magnitude of their regression slope (purple: steeper slopes, yellow: shallower slopes).

Viewing body shape differentiation in *Pristurus* in phylomorphospace (Figure 5) revealed broad overlap among habitat groups, though arboreal (tree-dwelling) species were somewhat more separated in morphospace. Rock-dwelling species occupied a slightly larger region of morphospace as compared with the other groups, though this pattern was not statistically significant (Supplementary Material). Intriguingly, when viewed in relation to body size, large *Pristurus* species were not localized to a particular region of morphospace, nor were smaller species. Instead, the largest rock-dwelling species were found in close proximity to the smallest ground-dwelling species, indicating that they were similar in overall body shape. Likewise, the smallest rock-dwelling species were found close to large ground-dwelling species in morphospace, indicating they displayed similar body shapes as well.

**Figure 5.**
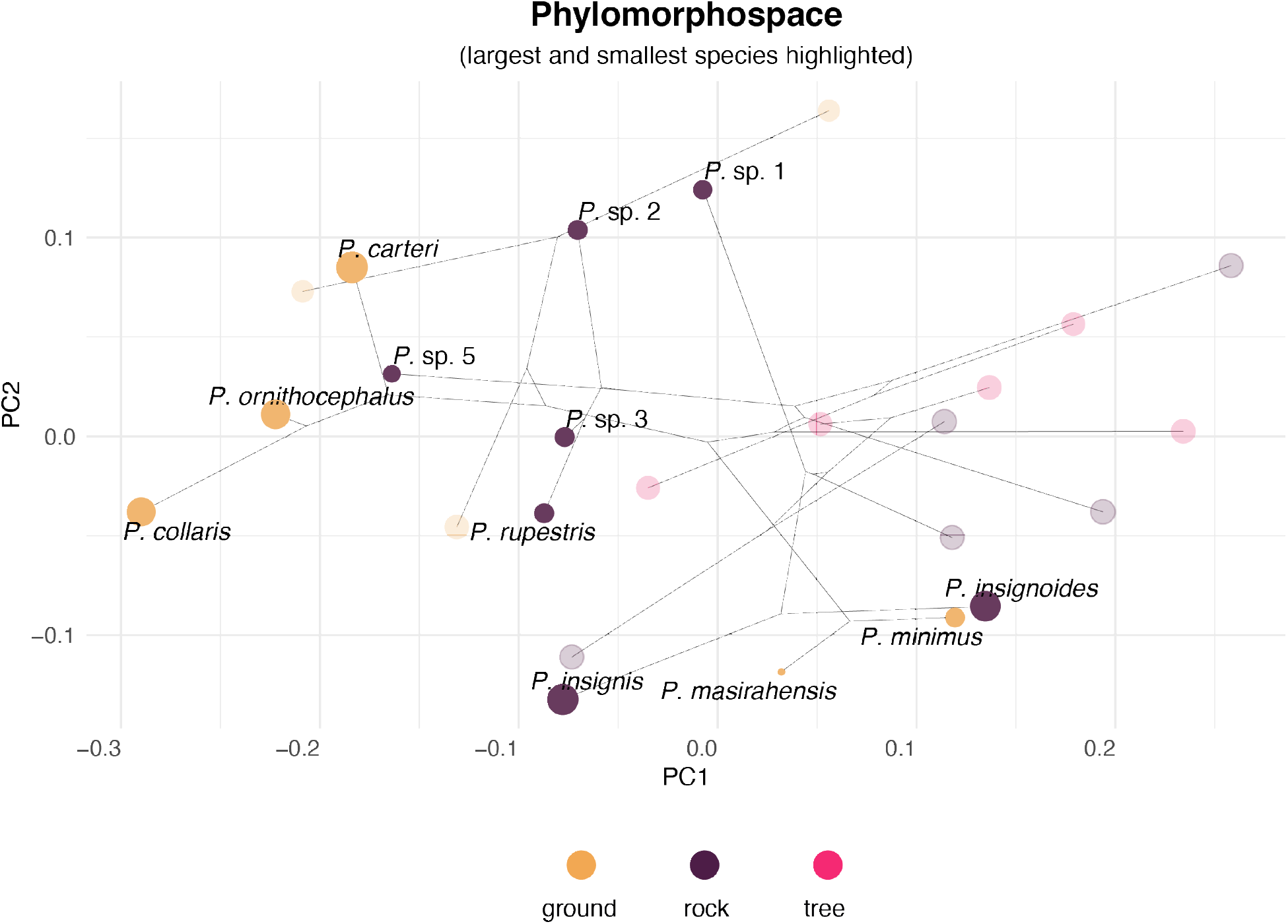
Phylomorphospace of *Pristurus*, based on residuals from a phylogenetic regression of body measurements on size (SVL). Species means are colored by habitat use: ground (beige), rock (dark purple), and tree (magenta). Large and small rock-dwelling and ground-dwelling are highlighted with darker colors to highlight their differentiation and relative positions in morphospace.

Finally, when representative specimens were scaled to a similar body size (Figure 6), the consequences of differences in allometric trends on body proportions became apparent. Here, larger ground-dwelling *Pristurus* species displayed disproportionately larger heads and limbs as compared with large *Pristurus* species utilizing other habitat types. Conversely, smaller rock-dwelling species were found to have disproportionately larger heads and limbs as compared with smaller ground-dwelling species. These patterns corresponded closely with those identified in morphospace (Figure 5), where large ground-dwelling species were similar in body form to small rock-dwelling species, while small ground-dwelling species were similar in body form to large rock-dwelling species (Figure 6). Thus, synthesizing the patterns revealed in the phylomorphospace with those from the other analyses revealed that the same body shape could be obtained in different ways, as determined by subtle differences in allometric slope across habitats, combined with body size differences. As such, species with similar body shapes displayed differing overall size, were found in distinct habitats, and exhibited different allometric trends.

**Figure 6.**
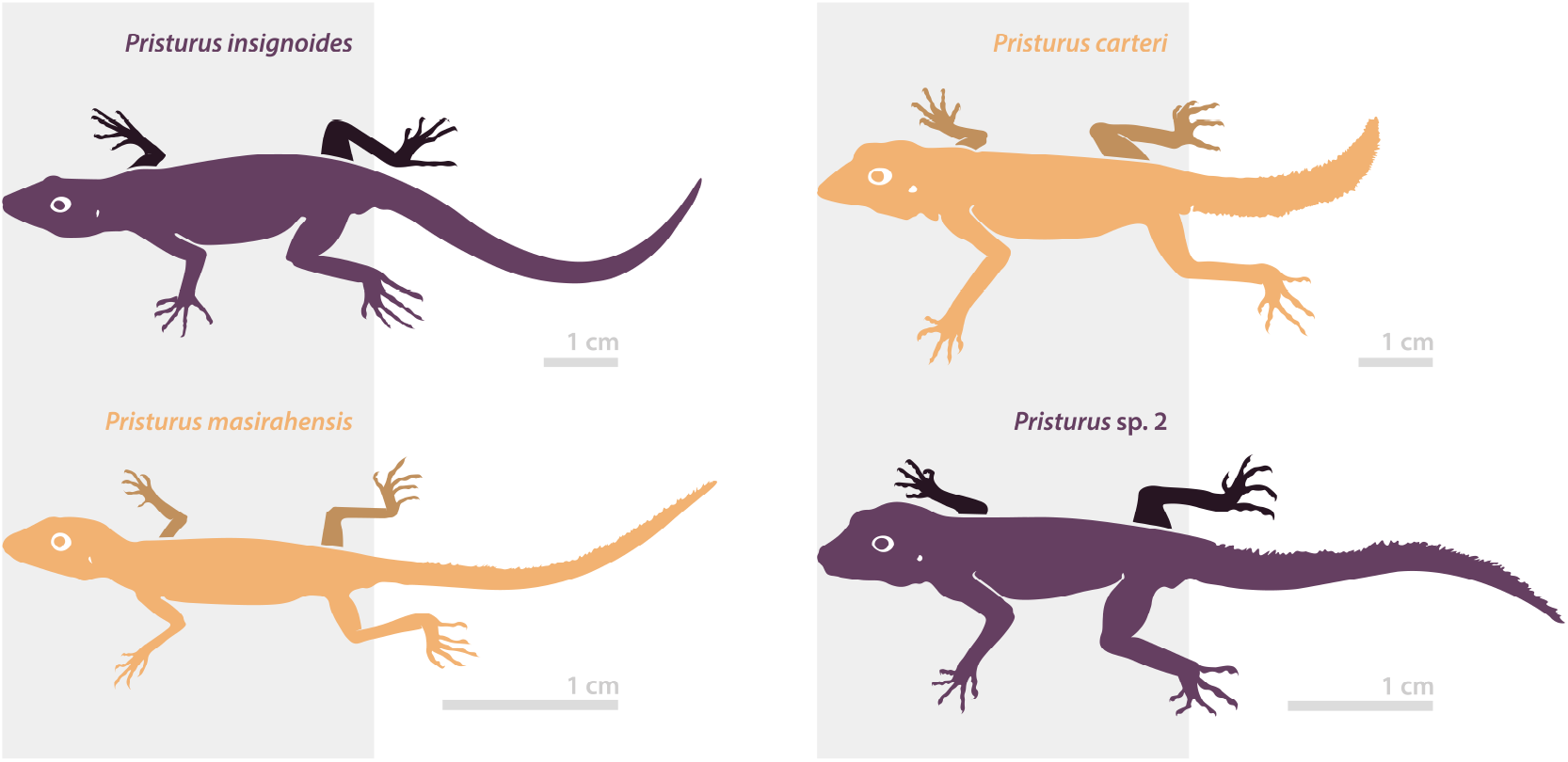
Representative specimens (based on real specimens) from large and small *Pristurus* species, colored by habitat use: ground (beige) and rock (dark purple). Specimens are scaled to a common body size (SVL, gray rectangles) to emphasize the relative differences in limb and head proportions. Relatively slender-headed and short-limbed species shown on the left. Original scale shown as the gray bar.

## 4. Discussion

Elucidating the selective forces that generate patterns of phenotypic diversity is a major goal in evolutionary biology. For species that utilize distinct habitats, disentangling the causes of phenotypic differentiation across those habitats is essential for our understanding of how natural selection operates and how evolution proceeds. In this study, we evaluated the role of potential drivers of body shape differentiation in the geckos of the genus *Pristurus*. To this end, we compared allometric trends and levels of integration among *Pristurus* occupying distinct habitats, interrogated allometric patterns at both the static and evolutionary levels, and related these trends to diversification in body form. Our findings have several important implications for how ecological specialization, phenotypic integration, and body form evolution along allometric trajectories relate to patterns of phenotypic diversity generally, and the evolution of phenotypic diversification in *Pristurus* in particular.

First, our analyses revealed that patterns of body shape allometry and morphological integration are relatively distinct in ground-dwelling *Pristurus* lizards, as compared with *Pristurus* occupying other habitats. Specifically, we found that multivariate vectors of regression coefficients differed significantly from what was expected under isometry (Table 2) for taxa utilizing all habitat types (ground, rock, tree), indicating that in *Pristurus*, allometric scaling patterns predominate. Further, our interrogation of allometric trends revealed differences between habitat types, where ground-dwelling *Pristurus* displayed steeper (i.e., positively allometric) trends for both head and limb traits, while rock and tree-dwelling taxa displayed shallower (negatively allometric) trends for head traits and more varied patterns for limb proportions. Biologically, these patterns revealed that not only does shape differ between large and small *Pristurus*, but this pattern differs across habitat types. Specifically, large ground-dwelling *Pristurus* present disproportionately larger heads and longer limbs relative to large individuals in other habitats, while small ground-dwelling *Pristurus* exhibit disproportionately smaller heads and shorter limbs (Figure 3). These findings are consistent with previous work at the macroevolutionary level [30], where large ground species were also found to display disproportionately large heads and long limbs.

Second, our findings revealed that rock-dwelling *Pristurus* show a converse pattern, where smaller individuals displayed relatively larger heads, while larger individuals have smaller heads relative to their body size. These allometric patterns also corresponded with findings at macroevolutionary scales [30], where similar patterns at the species level were observed. Regarding relative limb proportions, we found a high variability among small rock-dwelling species rather than a common pattern (Figure 3B). Indeed, earlier work in the subclade comprising several of these species (the *P. rupestris* species complex) found two well-differentiated phenotypes in populations of these lineages segregated by elevation [51]. These two ecotypes, defined as ‘slender’ and ‘robust’, differed in their head and limb characteristics. Our work is consistent with this, and extends these patterns to the allometric realm. Tejero-Cicuéndez et al. [30] also performed habitat ancestral estimation, finding that the rock habitat was the most likely ancestral condition in the group, with subsequent colonization by *Pristurus* of ground habitats. When patterns of allometry are viewed through this lens, it suggests the hypothesis that habitat shifts from rock-dwelling to ground-dwelling incurred a concomitant evolutionary shift in allometric trajectories as well [39]. Indeed, our analyses are consistent with this hypothesis, as allometric trends are inferred to be more rock-like towards the root of the *Pristurus* phylogeny (Figure 3), with subsequent shifts along branches leading to ground-dwelling species. This further suggests that the segregation in body size and shape through differential allometric relationships across habitats responds to adaptive dynamics concerning the colonization of new habitats. Thus, in *Pristurus*, there is support for the hypothesis that colonization of ground habitats has been a trigger for morphological change [30], as there appears to be a link between shifts in allometric trajectories as a result of habitat-induced selection, and differential patterns of body shape observed across taxa. More broadly, these findings are consistent with prior discoveries in other lizards, where the differential selective pressures imposed by rocky and ground habitats have resulted in the differentiation of head and limb morphology [43,51–53]. Indeed, such phenotypic differences resulting from the effects of habitat-based ecological selection have been extensively documented in reptiles as well as in other vertebrates [9,54–60], and our work in *Pristurus* thus contributes to this growing body of literature.

Another important finding of our study was the strong concordance between static allometry across individuals and evolutionary allometry among *Pristurus* species. Our analyses revealed small pairwise angles between static and evolutionary allometry vectors, indicating that allometric trends at these two hierarchical levels were oriented in similar directions and were essentially parallel. As such, size-associated changes in body shape among individuals were predictive of evolutionary shifts across taxa at higher macroevolutionary scales. This in turn, suggests that body shape evolution in *Pristurus* follows an allometric line of least resistance [61]. In other empirical systems, a similarly tight correspondence between static and evolutionary allometry has also been observed [61–65], though the trend is not universal across all taxa or traits [19,66]. Nonetheless, when such trends are present, they imply that allometric trajectories impose a prevailing influence on the magnitude, direction, and rate of phenotypic change across the phylogeny. Our work in *Pristurus* contributes to the growing literature on this topic, and suggests that perhaps such patterns may be more widespread.

Given the observation that static and evolutionary allometry in *Pristurus* are so concordant, an obvious question is: why might this be the case? One possible explanation is that when genetic covariation remains relatively constant, selection on body size will generate an evolutionary allometric trajectory along the trend described by static allometry [67,68]. Here, allometry effectively acts as a constraint on evolutionary change, as size-associated shape changes at one hierarchical level are linked to changes at another level [63,66,69]. Further, when this is the case, one may also expect high levels of phenotypic integration in traits associated with body size changes. Indeed, our analyses reveal precisely this pattern in *Pristurus*, with the highest levels of integration in the group (ground-dwelling) whose static allometry is most similar to that of evolutionary allometry. Thus, our results reveal that patterns of trait covariation are more constrained in ground-dwelling species, such that their differences in body form are most likely found along the primary allometric axis. When viewed in this light, integration and allometry may thus be interpreted as potential drivers that facilitate morphological change, as they provide a phenotypic pathway through adaptive lines of least resistance that enable rapid evolutionary changes in particular phenotypic directions but not in others [22,27]. The fact that ground-dwelling species in *Pristurus* have been found to have the widest phenotypic disparity, greatest range of body sizes, and highest rates of morphological evolution [30] are all consistent with this hypothesis, and suggest that in this group, integration describes the path of morphological evolution along allometric lines of least resistance.

Finally, interpreting the observed patterns of phenotypic integration and allometry relative to habitat-specific differences helps to shed light on the possible pathways by which phenotypic diversity in *Pristurus* has evolved. For instance, prior work on this system [30] revealed that the colonization of new ecological habitats elicited strong ecological selection and phenotypic responses. This was particularly true of the invasion of ground habitats, where ground-dwelling species displayed the largest variation in body size in the genus. This observation implies some level of ecological selection on body size. In lizards, the ecological context in which species exist is known to play a pervasive role in body size evolution [70–72], as it does in other animal groups [73–77]. While to date this has not been thoroughly explored in *Pristurus*, the evolutionary patterns revealed by our analyses suggest that the body size diversity in this clade conforms, at least in part, with patterns expected under ecological selection on body size. Intriguingly, such patterns are not only observed in ground- and rock-dwelling taxa, but also in arboreal species, whose restricted phenotypic diversity in both size and shape (Figures 3 & 5) is consistent with strong ecological selection in the arboreal habit [78,79]. Furthermore, our study identified the presence of strong integration and allometric trajectories, such that evolutionary changes in body size elicit corresponding changes in body shape. However, these trends differed significantly across habitats, implying that, at evolutionary scales, these trends serve to channel phenotypic responses to selection, but do so in differing directions for the different habitat groups. This, in turn, suggests that *Pristurus* species occupying different habitats display differing combinations of body size with body shape. The evolutionary consequence of ecological selection is that species have evolved similar shapes (Figure 6), but do so in differing habitats, and at different body sizes (Figure 5). Therefore, the phenotypic diversity observed in *Pristurus* is best explained as the result of a complex interplay between ecological selection, body size differentiation, and differing allometric trajectories across ecological habitats.

## Acknowledgments

We are very grateful to J. Roca, M. Metallinou, K. Tamar, J. Šmíd, R. Vasconcelos, R. Sindaco, F. Amat, Ph. de Pous, L. Machado, J. Garcia-Porta, J. Els, T. Mazuch, T. Papenfuss, and all the people from the Environment Authority, Oman, for their help in different aspects of the work.

## Data availability statement

All the data used in this study are available on DRYAD from a previous study: https://doi.org/10.5061/dryad.xwdbrv1f6 [32]. The scripts for implementing all analyses and generating the figures in this manuscript can be found in the Supplementary Material and in a GitHub repository (and on DRYAD upon acceptance).

