## Supplementary Material for "Evolution along allometric lines of least resistance: Morphological differentiation in *Pristurus* geckos"

#### 1: R-script for Article Computations and Flow of Allometry Operations

Below is an R-script that may be used to reproduce all statistical analyses found in the paper. The data are found on DRYAD: (doi:10.5061/dryad.xwdbrv1f6 [1]). Additional scripts used to generate publication-ready plots, and scripts for the additional plots, are found below.

```
libs <- c('geomorph', 'RRPP', 'phytools', 'geiger', 'tidyverse')
easypackages::libraries(libs)

# 0: Data Prep
data0 <- read.table('Analyses/data/morpho/morpho_pristurus.csv', sep = ';',
               dec = '.', header = TRUE, stringsAsFactors = TRUE)
  sp.to.keep <- names(which(table(data0$species) >= 5))
data <- data0[data0$species %in% sp.to.keep, ]
  data$species <- droplevels(data$species)
  data$SVL <- log(data$SVL)
shape <- as.matrix(log(data[, 8:ncol(data)]))
rdf <- rrpp.data.frame(svl = data$SVL, shape = shape,
           habitat = data$habitat_broad, species = data$species)
tree0 <- read.nexus('Analyses/data/phylogeny/pristurus_tree_final.nex')
LS.mns <- pairwise(lm.rrpp(shape~species, data = rdf, iter=0),
                  groups = rdf$species)$LS.means[[1]]
sz.mn <- tapply(rdf$svl,rdf$species,mean)
hab.mn <- as.factor(by(rdf$habitat,rdf$species,unique))
levels(hab.mn) <- levels(rdf$habitat)
tree <- treedata(phy = tree0, data = LS.mns)$phy
C <- vcv.phylo(tree)

# 1: Evolutionary Allometry
allom.sp <- lm.rrpp(LS.mns~sz.mn, Cov = C)
allom.ind <- lm.rrpp(shape~svl, data = rdf)
anova(allom.sp)
anova(allom.ind)

M <-rbind(coef.sp <- allom.sp$LM$gls.coefficients[2,],
        coef.ind <- allom.ind$LM$coefficients[2,])

acos(RRPP:::vec.cor.matrix(M))*180/pi
```

```
# 2: Comparison of multivariate allometry among habitat types
fit.hab <- lm.rrpp(shape~svl*habitat, data = rdf)
  anova(fit.hab)

# 2A: Compare habitat vectors versus isometry and to each other
  #H_0: isometry as common slope model
mn.sz <- tapply(rdf$svl,rdf$habitat,mean)
mn.shape <- rowsum(rdf$shape, rdf$habitat)/as.vector(table(rdf$habitat))
coef.iso <- c(1,1,1,1,1,1,1,1)
intercepts <- mn.shape - t(tcrossprod(coef.iso,mn.sz))
X <- model.matrix(~rdf$svl+rdf$habitat)
b <- rbind(intercepts[1,],coef.iso,intercepts[2,]-intercepts[1,],
           intercepts[3,]-intercepts[1,])
preds <- X%*%b
E.iso <- rdf$shape - preds
perms <- RRPP:::perm.index(n = fit.hab$LM$n, iter = 999)
slopes <- list()
for(j in 1:1000){
  slopes[[j]] <- pairwise(lm.rrpp((preds+E.iso[perms[[j]],]) ~ rdf$svl*rdf$habitat,
          iter=0), groups = rdf$habitat,covariate = rdf$svl)$slopes[[1]]
}
slp.ang <- lapply(1:1000, function(j)
     acos(RRPP:::vec.cor.matrix(rbind(slopes[[j]],coef.iso)))*180/pi)

slp.hab.obs <- slp.ang[[1]]
slp.Z <- RRPP:::effect.list(slp.ang)
slp.P <- RRPP:::Pval.list(slp.ang)

slp.hab.obs
slp.Z
slp.P

# 2B: Compare evolutionary and static (habitat) allometry
  #H_0: common slope isometry
slp.ang.ev <- lapply(1:1000, function(j) cos(RRPP:::vec.cor.matrix(rbind(coef.evol,
              slopes[[j]])))*180/pi)

slp.hab.ev.obs <- slp.ang.ev[[1]]
slp.Z.ev <- RRPP:::effect.list(slp.ang.ev)
slp.P.ev <- RRPP:::Pval.list(slp.ang.ev)

slp.hab.ev.obs
slp.Z.ev
slp.P.ev

res <- cbind(slp.hab.ev.obs[-1,1],slp.Z.ev[-1,1],slp.P.ev[-1,1])
colnames(res) <- c("Angle","Effect Size", "P-value")
```

```r
rownames(res) <- c("Ev vs. Ground", "Ev vs. Rock", "Ev vs. Tree")
res

# 3: Map allometry slopes on phylogeny
head.scores <- two.b.pls(shape[, c(2:4)], rdf$svl)$XScores[, 1]
limb.scores <- two.b.pls(shape[, 5:8], rdf$svl)$XScores[, 1]

coef.head <- lm.rrpp(head.scores ~ rdf$svl*rdf$species)$LM$coefficients
coef.limb <- lm.rrpp(limb.scores ~ rdf$svl*rdf$species)$LM$coefficients

head.slp <- coef.head[grep('svl', rownames(coef.head)), ]
  head.slp[-1] <- head.slp[-1] + head.slp[1]
limb.slp <- coef.limb[grep('svl', rownames(coef.limb)), ]
  limb.slp[-1] <- limb.slp[-1] + limb.slp[1]
names(limb.slp) <- names(head.slp) <- levels(rdf$species)

cor(head.slp,limb.slp)
plot(head.slp,limb.slp)

contMap(tree = tree, x = head.slp, outline = FALSE)
cm.limb <- contMap(tree = tree, x = limb.slp, outline = FALSE)

# 4: Compare Integration
lindims.gp <- lapply( split( shape[,1:ncol(shape)], rdf$habitat),
                      matrix, ncol=ncol(shape))
Vrel.gp <- Map(function(x) integration.Vrel(x), lindims.gp)
c(Vrel.gp$ground$ZR,Vrel.gp$rock$ZR,Vrel.gp$tree$ZR)
out <- compare.ZVrel(Vrel.gp$ground, Vrel.gp$rock, Vrel.gp$tree)
summary(out)

# 5: phylomorphospace of size-standardized data (residuals)
shape.res <- residuals(allom.sp)
pca.w.phylo <- gm.prcomp(shape.res, phy = tree)
plot(pca.w.phylo, phylo = TRUE, pch = 21, bg = 'black',
     phylo.par = list(node.labels = FALSE))
```

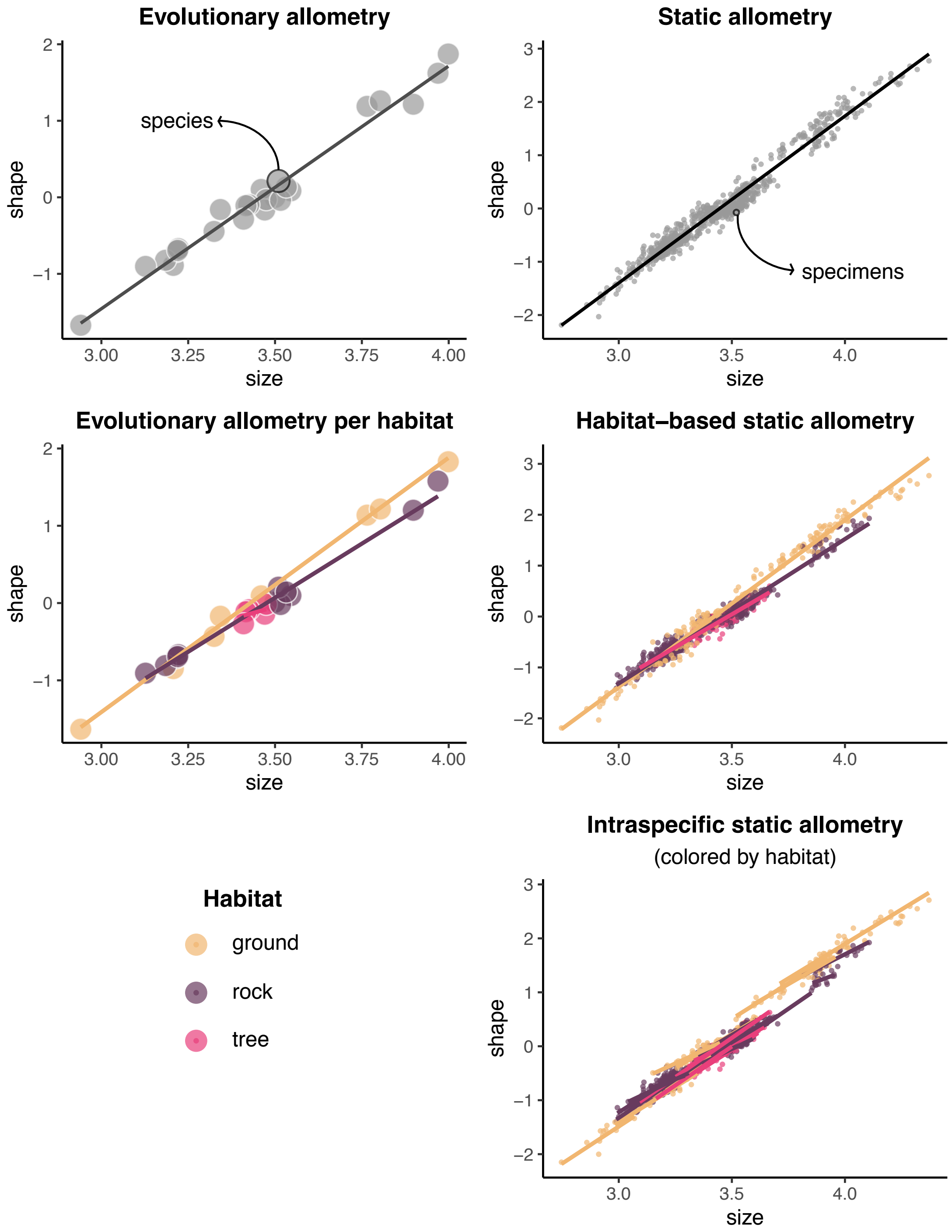


Figure 1: Visual depiction of flow of operations for interrogating levels of allometry. Colors designate the different habitat groups.

### 2: Additional Analyses and Visualizations

Here we provide additional analyses which complement those found in the article.

##### Inspection of Regression Coefficients (Slopes)

Here are the regression slopes for each habitat group, found from our linear model. These display differences in allometry among groups, variable by variable.

```
## Warning in treedata(phy = tree0, data = LS.mns): The following tips were not found in 'data
##  Pristurus_adrarensis
##  Pristurus_flavipunctatus
##  Pristurus_sp12
##  Pristurus_sp4
##  Pristurus_sp9
```

```
fit.hab <- lm.rrpp(shape~svl*habitat, data = rdf)
pw.hab1 <- pairwise(fit.hab, groups = rdf$habitat, covariate = rdf$svl)
  slp.hab <- pw.hab1$slopes[[1]] #slopes by habitat
slp.hab
```

```
##             TrL        HL        HW        HH      Lhu       Lun       Lfe
## ground 1.027941 1.0283300 1.1933370 1.0678123 1.185343 1.2736867 1.3186471
## rock   1.096908 0.9409198 0.8131810 0.8746610 1.044875 1.1121324 1.0553304
## tree   1.086165 0.8087240 0.7954059 0.7428799 1.033895 0.9998545 0.9079136
##              Ltb
## ground 1.1584966
## rock   1.0663836
## tree   0.9311451
```

#### Traitgrams of Individual Trait Allometry

In the main article we provided traitgrams of allometric slopes for composites of head traits (Figure 3A) and limb traits (Figure 3B). Here we provide traitgrams for the allometric relationship of each body trait separately. As in the main article, traitgrams are visualized from an evolutionary mapping of body size (SVL), and the color represents changes in the allometric slope for each phenotypic trait, found from an evolutionary mapping of the species-level slopes under a Brownian motion model of evolution.

Here we see that there are differential allometric dynamics for some morphological variables. Most notably, large species inhabiting rocky habitats show an increase in the allometric slope for the length of the tibia (Ltb), while the rest of the limb segments (Lhu, Lun, and Lfe) have a decreasing trend. Conversely, these species show virtually identical trends for the head shape variables (HL, HW, and HH), where the size increase has occurred together with a reduction of the allometric slope. Overall, large ground-dwelling species show opposite allometric trends to those of the large rock dwellers in most variables. However, it might be interesting to notice that one of these large ground species, *Pristurus ornithocephalus* (the fourth largest species of the genus, and the second largest ground species) presents unique allometric tendencies relative to the other large ground species, both for all head variables and some of the limb variables (e.g., Lhu and Lfe). This might be reflecting species-specific ecological dynamics, and more detailed data and analyses (e.g., geometric morphometrics) could shed light on this morphological pattern.

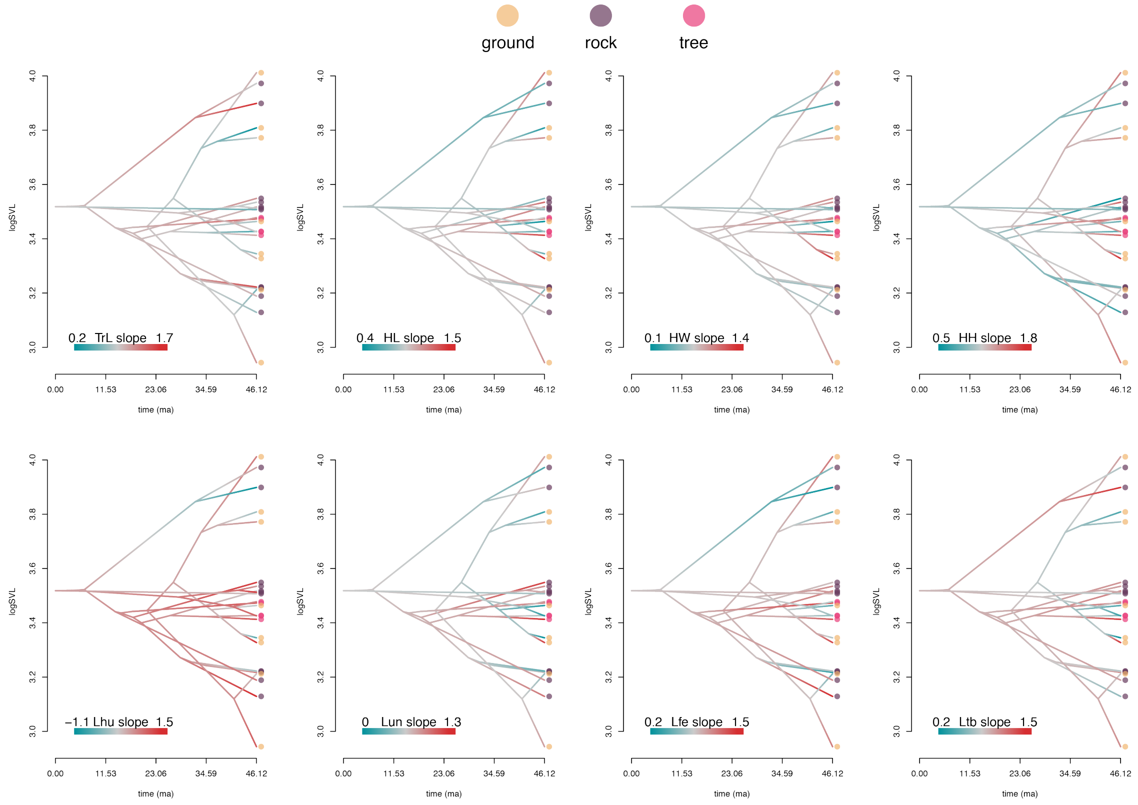


Figure 2: Traitgrams of regression slopes for each phenotypic variable. Colors at the tips designate habitat groups.

#### Evolutionary Mapping of Head & Limb Allometry

In the main article, allometric trends in both head dimensions were mapped onto the phylogeny under a Brownian motion model of evolution to discern macroevolutionary changes across the phylogeny. These were visualized on traitgrams, where body size differences were optimized (main article: Fig. 3). Here we present evolutionary mappings of allometric trends individually, so that increases and decreases in allometric slopes across the phylogeny are more readily interpreted. A summary of these patterns was described in the main article.

Briefly, these plots show that changes in allometry were not concentrated to particular regions of the phylogeny, but rather displayed both increases and decreases in allometry of both the head traits and the limb traits occurred repeatedly in this group.

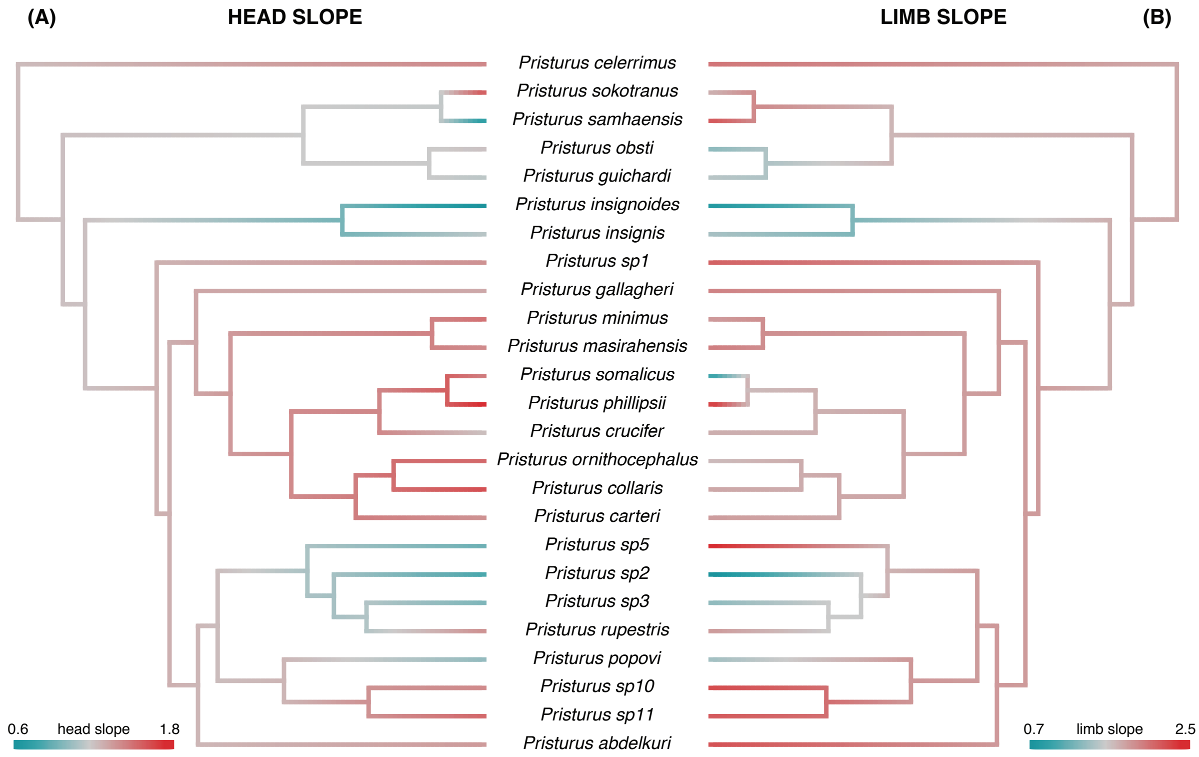


Figure 3: Evolutionary mapping of regression slopes describing the relationship of (A) head morphology versus body size, and (B) limb proportions versus body size.

##### Morphological Integration

In the main article, we performed analyses of morphological integration on the set of body traits representing body form. Here we perform the same analysis, using size-standardized data.

```
shape.2 <- shape - rdf$svl
shape.gp <- lapply( split( shape.2[,1:ncol(shape.2)], rdf$habitat),
                    matrix, ncol=ncol(shape.2))
Vrel.gp.shp <- Map(function(x) integration.Vrel(x), shape.gp)
c(Vrel.gp.shp$ground$ZR,Vrel.gp.shp$rock$ZR,Vrel.gp.shp$tree$ZR)
```

```
## [1] 2.403052 1.904412 1.224868
```

```
out.shp <- compare.ZVrel(Vrel.gp.shp$ground, Vrel.gp.shp$rock, Vrel.gp.shp$tree)
summary(out.shp)
```

```
##
## Effect sizes
##
## Vrel.gp.shp$ground   Vrel.gp.shp$rock   Vrel.gp.shp$tree
##         -0.1468813         -0.9606380         -0.9189899
##
## Effect sizes for pairwise differences in rel.eig effect size
##
##                    Vrel.gp.shp$ground Vrel.gp.shp$rock Vrel.gp.shp$tree
## Vrel.gp.shp$ground           0.000000        9.3926169        6.1015621
## Vrel.gp.shp$rock             9.392617        0.0000000        0.3556259
## Vrel.gp.shp$tree             6.101562        0.3556259        0.0000000
##
## P-values
##
##                    Vrel.gp.shp$ground Vrel.gp.shp$rock Vrel.gp.shp$tree
## Vrel.gp.shp$ground       1.000000e+00     5.852770e-21     1.050368e-09
## Vrel.gp.shp$rock         5.852770e-21     1.000000e+00     7.221208e-01
## Vrel.gp.shp$tree         1.050368e-09     7.221208e-01     1.000000e+00
```

##### Phylomorphospace and Disparity

In the main article we presented a representation of phylomorphospace using size-standardized shape variables. Here we present a test of disparity of the size-standardized data. Additionally, we present a similar plot for the unadjusted species means. In addition, we calculate the phenotypic disparity in among species in each habitat group, and compare these using permutation.

First, we show that in the size-standardized morphospace, morphological disparity does not differ across habitat groups. There are no differences.

```
fit.lm <- lm.rrpp(shape.res~hab.mn, Cov =C)
PW.lm <- pairwise(fit.lm,groups = hab.mn)
summary(PW.lm, test.type = "var")
```

```
##
```

```
## Pairwise comparisons
##
## Groups: ground rock tree
##
## RRPP: 1000 permutations
##
##
## Observed variances by group
##
##    ground      rock      tree
## 0.2938297 0.3015873 0.2260537
##
## Pairwise distances between variances, plus statistics
##                       d UCL (95%)          Z Pr > d
## ground:rock 0.007757614 0.1477325 -1.4732960  0.921
## ground:tree 0.067776058 0.1503602  0.2855938  0.421
## rock:tree   0.075533672 0.1020809  1.0336714  0.158
```

Next we generate a phylomorphospace for the unadusted species means. Not surprisingly, PC1 of the morphospace represents size, and in fact scores on PC1 are highly correlated with SVL ($\rho = 0.987$).

```
pca.w.phylo2 <- gm.prcomp(LS.mns, phy = tree)
cor(sz.mn,-1*pca.w.phylo2$x[,1])
```

```
## [1] 0.9879806
```

```
plot(pca.w.phylo2, phylo = TRUE, pch = 21, bg = 'black',
     phylo.par = list(node.labels = FALSE))
```

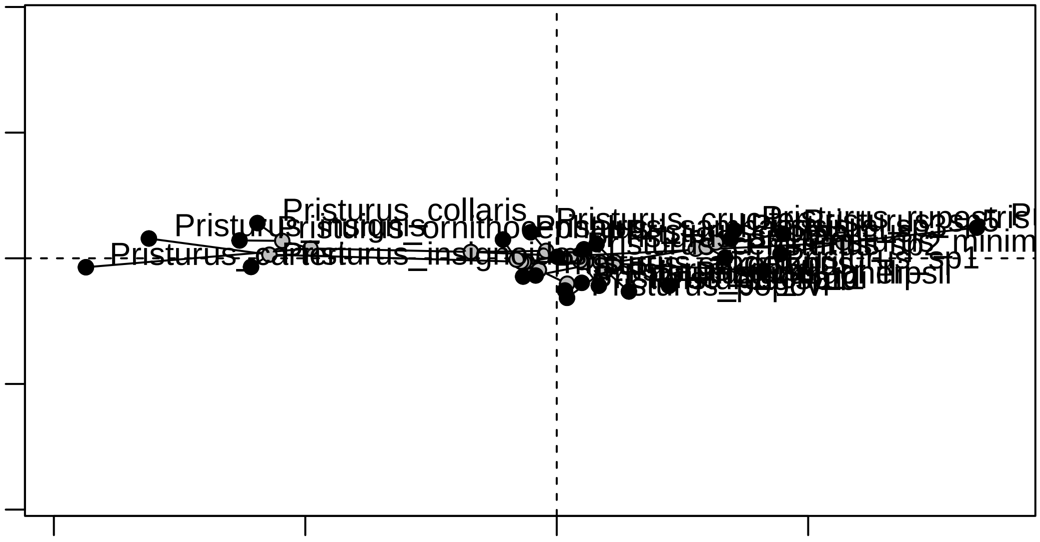


Disparity was estimated from a phylogenetic anova, obtained using RRPP. Here we observed that the ground-dwelling species display greater disparity then do the other two habitat groups.

```
fit.lm <- lm.rrpp(LS.mns~hab.mn, Cov =C)
PW.lm <- pairwise(fit.lm,groups = hab.mn)
summary(PW.lm, test.type = "var")
```

```
##
## Pairwise comparisons
##
## Groups: ground rock tree
##
## RRPP: 1000 permutations
##
##
## Observed variances by group
##
##    ground      rock      tree
## 18.385078  7.666641  5.635931
##
## Pairwise distances between variances, plus statistics
##                    d UCL (95%)           Z Pr > d
## ground:rock 10.71844  7.924989  2.12714952  0.013
## ground:tree 12.74915  8.616215  2.41223861  0.002
## rock:tree    2.03071  5.745950 -0.03422601  0.521
```
